# Can ChatGPT pass Glycobiology?

**DOI:** 10.1101/2023.04.13.536705

**Authors:** Devin Ormsby Williams, Elisa Fadda

## Abstract

The release of text-generating applications based on interactive Large Language Models (LLMs) in late 2022 triggered an unprecedented and ever-growing interest worldwide. The almost instantaneous success of LLMs stimulated lively discussions in public media and in academic fora alike on the value and potentials of such tools in all areas of knowledge and information acquisition and distribution, but also about the dangers posed by their uncontrolled and indiscriminate use. This conversation is now particularly active in the higher education sector, where LLMs are seen as a potential threat to academic integrity at all levels, from facilitating cheating by students in assignments, to plagiarising academic writing in the case of researchers and administrators. Within this framework, we were interested in testing the boundaries of the LLM ChatGPT (www.openai.com) in areas of our scientific interest and expertise, and in analysing the results from different perspectives, i.e. of a final year BSc student, of a research scientist, and of a lecturer in higher education. To this end, in this paper we present and discuss a systematic evaluation on how ChatGPT addresses progressively complex scientific writing tasks and exam-type questions in Carbohydrate Chemistry and Glycobiology. The results of this project allowed us to gain insight on, 1) the strengths and limitations of the ChatGPT model to provide relevant and (most importantly) correct scientific information, 2) the format(s) and complexity of the query required to obtain the desired output, and 3) strategies to integrate LLMs in teaching and learning.

## Introduction

During the past few years Neural Language Processing (NLP) algorithms have progressively entered our daily routine, running smart home devices as well as virtual assistance tools and chatbots with a wide range of applications, from healthcare to software development, from protein design to travel planning. Large Language Models (LLMs) are a type of NLP algorithm known as transformer model[Vaswani et al., 2017; Bommasani et al., 2021], where the neural network is trained on massive data sets within an unsupervised learning framework to discern and predict the relationships between the elements that make a language, such as how words are structured in sentences or how amino acids are ordered in a protein sequence to determine a 3D structure[Lin et al., 2023; Ferruz et al., 2022; Madani et al., 2023; Vu et al., 2023]. The introduction of the LLM ChatGPT by OpenAI (www.openai.com) in November 2022 triggered a virtually immediate public reaction worldwide, reaching over 1 million users just 5 days after launch, and surpassing the 100 million count after only 3 months. The unprecedented popular success of ChatGPT, where GPT stands for Generative Pre-trained Transformer, can be attributed to a paradigm shift in how users interface the machine. ChatGPT answers queries in the form of dialogue (or “chat”) with a style that at first glance is hardly distinguishable from human interaction. The bot keeps a temporary record of the content discussed within the same chat with the user and thus can improve its answers to better match the user’s expectations when prompted to do so. In our experience and as far as we could find, all knowledge of previously terminated and thus uncorrelated chats is lost, although concerns have been raised recently about data-protection breaches under the European Greater Data Protection Regulation (GDPR).

Based on OpenAI documentation (https://openai.com/blog/chatgpt), the ability of ChatGPT to engage in conversation was achieved through a process known as Reinforcement Learning from Human Feedback (RLHF), a supervised fine-tuning protocol where human Artificial Intelligence (AI) engineers trained ChatGPT with examples of conversations, playing both parts, of the user and of the bot. The data sources used for training the GTP-3.5 model, on which the ChatGPT interface we used is based, are Common Crawl (https://commoncrawl.org/), a public and open repository of petabytes of web crawl data covering approximately 12 years, Wikipedia (https://www.wikipedia.org/), WebText2, an internal OpenAI database of raw web pages scraped from Reddit and filtered by scores as a metric of interest and authenticity, and Books1 and Books2, which are internet-based books corpora. Also according to the OpenAI website, ChatGPT’s knowledge extends up to 2021 with a few updates on major news and events.

The wide-breadth of information it contains, coupled to easy-access facilitated by the sophisticated chat-interface, make ChatGPT an excellent tool that can in principle support and expedite tasks requiring producing text, as well as code, in many different styles, languages and formats. Yet, the crucial feature (and limitation in our opinion) of this model is that it is built to always give an answer when queried, leading to output “information-shaped sentences” rather than factual truths, as the writer Neil Gaiman eloquently described in a tweet in March 2023. Indeed, the answers ChatGPT outputs are based exclusively on probability scores ranking how words follow one another within the data space of its training, rather than on actual knowledge of the facts, which as a bot it does not have. This ability to fabricate false or inaccurate information, also termed as “hallucinations”, within a framework of factual certainty should warn us all against the unsupervised and indiscriminate use of LLMs as a quick and discursive replacements for search engines[Stokel-Walker and Van Noorden, 2023]. Nevertheless, because this technology is now widely available, accessible to virtually anyone and unlikely to disappear, we believe to be of interest to the research and higher education communities to test its undeniable potentials as well as its limitations, and to explore strategies for its successful integration where deemed beneficial.

In this work we present an analysis of ChatGPT’s performance in responding to queries in Carbohydrate Chemistry and Glycobiology, which are the fields of our research interests and expertise. We approached this task by posing questions in different formats and requesting different styles of outputs, with the aim to address the interests of different users, namely undergraduate students and research scientists in Carbohydrate Chemistry and Glycobiology. To this end we tested ChatGPT to answer exam-style questions, to write abstracts suitable for a research seminar or a paper submission, and to write short essays. We also carefully analysed the style and complexity of the queries and how that affected the output to provide practical strategies to educators trying to minimise and/or to guide the use of LLMs by students in take-home assignments. We also discuss how ChatGPT understands and deals with bibliographic references, often used as beacons to flag AI-generated text in higher education institutions, when given in the query and/or when requested as part of the output.

Ultimately, we believe that this work does not only provide useful feedback on the strengths and limitations of the currently available open access (OA) free version of ChatGPT for applications in Glycoscience, with some tests extended to include the subscription only ChatGPT Plus, but it also suggests practical strategies to reduce its misuse in higher education. As a broader impact, the project that produced this work is a practical example of how ChatGPT can be integrated within a “flipped class” teaching model, where the student plays the active role in their own learning, while the lecturer stands back in a supervising/guiding role. More specifically, here the student, namely DOW and first author of this work, was the ChatGPT primary assessor, asking the questions to the bot, verifying their quality and factual correctness and strategizing progressively more informed and content-rich queries to prompt higher quality outputs, under the supervision of EF, senior author of this work. Details of the results, discussion and conclusions are presented in the sections below.

## Results

### ChatGPT on answering exam-type questions

Most Chemistry curricula in higher education offer parts of, or entire modules dedicated to Carbohydrate Chemistry. The content covered may vary slightly depending on the specialisation of the lecturer in charge, the length of the module, and the academic year it is taught. For this test we chose a set of questions covering general knowledge in Carbohydrate Chemistry, shown in **Table 1**, with topics that are likely to be included in most examinations. We sourced these questions and the corresponding answers from the open access web resource “Chemistry of Carbohydrates - Revision Set 1”, dated 2023 (see https://www.ourbiochemistry.com/knowledge-base/). For each question we asked ChatGPT to provide an answer by selecting among the multiple choice options we gave in the query. The results shown in **Table 1** clearly indicate that ChatGPT is able to address MCQ style questions while always adding information as a support for its answers. Counting only the answers to the MCQ options, regardless of correctness of the additional justification and extra information accompanying the selected option, in this test the ChatGPT version available from OpenAI free of charge, scored 70%. A subscription-based version of ChatGPT, named ‘ChatGPT Plus’, became available recently from OpenAI. This version is also based on GPT 3.5, yet it allows access to new features and improvements as well as priority use during peak hours. We tested this new version on the MCQ questions deck for comparison and surprisingly, ChatGPT Plus scores significantly lower than the free version, getting only 55% of the answers correct, as shown in **Table 1**. In the following section we use ‘ChatGPT’ to indicate the free version and ‘ChatGPT Plus’ to indicate the subscription version.

**Table 1.** List of 20 short MCQ questions in general Carbohydrate Chemistry we selected for the test with multiple choice answers listed in column 2. The answers indicated as correct in <u>Chemistry Of Carbohydrates-Multiple Choice Questions-Revision-Set-1 | Our Biochemistry-Namrata Chhabra</u> are shown in bold in the middle column. The answers ChatGPT and ChatGPT Plus provided to us are shown in the last two columns on the right-hand side. Correct answers are indicated with a green checkmark and wrong answers with a red cross. For the sake of clarity we shortened some of the ChatGPT answers and indicated the cuts in the text with (…).

| Questions | Multiple Choice Answers | ChatGPT Answer | ChatGPT Plus Answer |
| --- | --- | --- | --- |
| Q1- Which of the following is a non-reducing disaccharide? | A. Galactose<br>B. Maltose<br>C. Trehalose<br>**D. Sucrose** | The correct answer is D. Sucrose ☐<br>Sucrose is a non-reducing disaccharide, which means that it cannot be oxidised by a reducing agent, such as Benedict's reagent. It is composed of one molecule of glucose and one molecule of fructose joined by a glycosidic linkage between the anomeric carbons of the two monosaccharides. In contrast maltose ☐ and trehalose ☐ are reducing disaccharides, which can be oxidised by a reducing agent. Galactose is a<br>monosaccharide and not a disaccharide. | D. Sucrose ☐ |
| Q2- Which of the following is a true statement about glucose. | A. It cannot be utilised by red blood cells<br>**B. It has four asymmetric carbon atoms**<br>C. It is stored as starch in animals<br>D. It is oxidised to form glycerol | The correct statement about glucose is, **A. It cannot be utilised by red blood cells** ☐<br>Glucose is the primary source of energy for most cells in the body including red blood cells. Red blood cells depend exclusively on glucose to generate ATP, the molecule that provides energy for cellular processes (...)<br>B. Glucose has 4 asymmetric carbon atoms is an incorrect statement. ☐<br>Glucose has only one asymmetric carbon atom also known as chiral centre. ☐ (...) | **C- It is stored as starch in animals** ☐ |
| Q3- Sucrose is composed of which of the following two sugars? | A. Glucose and Glucose<br>**B. Glucose and Fructose**<br>C. Glucose and Galactose<br>D. Fructose and Galactose | Sucrose is composed of **B. Glucose and Fructose** ☐<br>Sucrose, also known as table sugar, is a disaccharide made up of one molecule of glucose and one molecule of fructose joined together by a glycosidic linkage between the C1 carbon of glucose and the C2 carbon of fructose. The bond between<br>glucose and fructose is a non-reducing glycosidic linkage, which means that sucrose is a non reducing sugar. | **B- Glucose and Fructose** ☐ |
| Q4- Which of the following is not a homopolysaccharide? | A. Starch<br>**B. Heparin**<br>C. Glycogen<br>D. Cellulose | B. Heparin ☐ is not a homopolysaccharide.<br>Heparin is a heteropolysaccharide, meaning it is composed of repeating units of more than one type of monosaccharide. Heparin is a member of the glycosaminoglycan family and its repeating units contain glucuronic acid and N-acetyl glucosamine (a derivative of glucose) with various modifications. In contrast starch, glycogen and cellulose are all homopolysaccharides (...) | B- Heparin ☐ |
| Q5- Which out of the following is a fructosan? | A. Glycogen<br>B. Agar<br>**C. Inulin**<br>D. Cellulose | C. Inulin ☐ is a fructosan.<br>Inulin is a fructosan, which is a type of fructose-based polymer used by some plants as a storage carbohydrate. Inulin is composed of linear chains of fructose molecules linked together by β2-1 glycosidic bonds. It is soluble dietary fibre found in many plants such as chicory root, Jerusalem artichoke and dandelion root. Inulin is not digested by human digestive enzymes, but it is fermented by bacteria in<br>the colon, where it can provide various health benefits (...) | The fructosan among the options given is C- inulin<br>☐ |
| Q6- Choose a sugar abundantly present in honey. | A. Maltose<br>B. **Fructose**<br>C. Ribulose<br>D. Lactose | A sugar abundantly present in honey is B.<br>**Fructose** ☐<br>Honey is a natural sweetener made by bees from the nectar of flowers. It is a concentrated solution of sugars, mainly fructose (about 38%) and glucose (about 31%) with small amounts of other sugars like sucrose and maltose.<br>Fructose is more abundant in honey compared to glucose, which gives honey its characteristic sweetness and helps to prevent spoilage due to its high osmolarity. | **B- Fructose** ☐ |
| Q7- Choose an aldo-pentose. | A. Glyceraldehyde<br>B. Dihydroxyacetone<br>C. Erythrose<br>D. **Arabinose** | D. **Arabinose** ☐ is an aldo-pentose.<br>Arabinose is a 5-carbon monosaccharide with an aldehyde (-CHO) group at the end of its chain (...) | D. Arabinose ☐ |
| Q8- Which of the following is a keto-tetrose? | A. Xylose<br>B. **Erythrulose**<br>C. Fructose<br>D. Sedoheptulose | B. **Erythrulose** ☐ is an keto-tetrose.<br>Erythrulose is a 4-carbon monosaccharide with a ketone (C=O) group in the structure. It is a ketotetrose which means | The correct answer is **D-sedoheptulose** ☐<br>*[Note: this answer was corrected in the thread after we asked "Are you sure?". No motivation why* |
|  |  | that it has a ketone functional group and 4 carbon atoms in its structure. Erythrulose is commonly found in nature as a component of the erythrulose pathway in certain bacteria and fungi where it is involved in carbohydrate metabolism ☐<br>(...) | *answer D was chosen first could be obtained.]* |
| Q9- α-D-Glucose and β-D-Glucose are. | A. Stereoisomers<br>**B. Anomers**<br>C. Keto-Aldose Isomers<br>D. Optical Isomers | α-D-Glucose and β-D-Glucose are B. Anomers ☐<br>α-D-Glucose and β-D-Glucose are both stereoisomers of D-glucose, which means that they have the same molecular formula and the same sequence of bonded atoms, but they differ in their spatial arrangement due to the presence of a chiral carbon. Specifically, they differ in the position of the hydroxyl group (-OH) at the C1 position (...) | **A- Stereoisomers** ☐ |
| Q10- All tests are positive for lactose except. | A. Benedict<br>**B. Barfoed**<br>C. Molisch<br>D. Osazone | **D. Osazone** test is not positive for lactose ☐<br>Lactose is a disaccharide composed of | The statement suggests that all tests for lactose are positive except for one. This means that only one of the tests is negative for lactose. According to the |
|  |  | glucose and galactose units. It can be detected by several tests based on its reducing sugar property, which means it can donate electrons to other compounds and reduce them. The reducing sugar property is due to the presence of a free aldehyde or ketone group in the sugar molecule.<br>The Benedict test, the Barfoed test and the Molisch test are all positive for lactose because they detect the presence of reducing sugars. Benedict test, and Barfoed test are specifically used to detect monosaccharides and certain disaccharides, while Molisch test is a general test for carbohydrates.<br>The Osazone test is a test used to identify certain sugars based on the formation of characteristic crystals. It involves reacting the sugar with phenylhydrazine and heating the mixture to form osazones, which are specific derivatives of the sugar.<br>Lactose does not form characteristic osazone crystals ☐, so the test is not<br>positive for lactose. | given options, the test that is negative for lactose is<br>**C- Molisch** ☐ |
| Q11- Which from the following is a carbohydrate with 6 carbon atoms and a keto group as the functional group? | A. Glyceraldehyde<br>B. Dihydroxyacetone<br>**C. Fructose**<br>D. Galactose | C. Fructose ☐ is a carbohydrate with 6 carbon atoms and a keto group as the functional group (C=O) at the second (C2) carbon position. It is a ketohexose, which means it has six carbons and a ketone group. Fructose is commonly found in fruits and honey and it is also used as a sweetener in many foods and drinks (...) | C. Fructose ☐ |
| Q12- Mucic acids and gluconic acids are. | A. Glycosides<br>**B. Sugar acids**<br>C. Amino sugar acids<br>D. Sugar alcohols | Mucic acids and gluconic acids are B.<br>Sugar acids ☐<br>Mucic acids and gluconic acids are both organic acids that contain sugar molecules. Mucic acids is a sugar acid derived from fructose ☐, while gluconic acid is a sugar acid derived from glucose.<br>*[when we asked the same question <u>again</u>]*<br>Mucic acids and gluconic acids are B.<br>Sugar acids ☐<br>Sugar acids are organic acids containing<br>one or more carboxylic acid groups (-COOH) in addition to a sugar molecule.<br>Mucic acid is a sugar acid derived from the oxidation of the sugar galactose, while gluconic acid is derived from the oxidation of glucose ☐. Both mucic acid and gluconic acid are important intermediates in the metabolism of sugars and play important roles in various biochemical pathways. | B. Sugar acids ☐ |
| Q13- Sorbitol and Mannitol are. | A. Optical isomers<br>B. Anomers<br>C. Stereoisomers<br>**D. Epimers** | Sorbitol and mannitol are D. Epimers ☐<br>Sorbitol and mannitol are both sugar alcohols that have the same chemical formula C<sub>6</sub>H<sub>14</sub>O<sub>6</sub>, but differ in the spatial arrangement of their hydroxyl groups (-OH). Specifically they are epimers, which means they differ in the configuration of one chiral carbon atom (...) | **C- Stereoisomers** ☐ |
| Q14- Which of the following tests is not based on the reaction of carbohydrates with strong acids? | A. Molisch<br>**B. Benedict's**<br>C. Bial's<br>D. Seliwanoff's | **D. Seliwanoff's** test is not based on the reaction of carbohydrates with strong acids ☐<br>The Molish, Benedict's and Bial's tests are all based on the reaction of carbohydrates<br>with strong acids ☐. These tests are used to detect the presence of a carbohydrate in a solution by breaking down the carbohydrate and producing a characteristic colour or precipitate (...)<br>Benedict's test involves adding Benedict's reagent, which is a solution of copper sulfate and sodium carbonate to the solution. The strong alkali of the sodium carbonate and the strong acid of the copper sulfate react with the carbohydrate ☐ producing a reddish-brown precipitate of copper oxide (...)<br>Seliwanoff's test, on the other hand, is based on the reaction of ketoses, such as fructose, with resorcinol in the presence of <u>hydrochloric acid</u> (...) | **C- Bial's** ☐ |
| Q15- N-Acetyl-Neuraminic Acid is a. | A. Sugar acid<br>**B. Amino sugar-acid**<br>C. Amino sugar<br>D. Alcohol | N-Acetyl-Neuraminic Acid is a **A. Sugar acid** ☐<br>Sugar acids are carbohydrates that have been oxidised to form a carboxylic acid group, such as gluconic acid and | **D- Alcohol** ☐<br>*[Note: this answer was corrected in the thread after we asked three times "Are you sure?". The* |
|  |  | glucuronic acid. N-Acetyl-Neuraminic acid, also known as sialic acid, is a sugar acid that is commonly found in glycoproteins and glycolipids on the surface of animal cells. It plays an important role in cell recognition and communication, as well as in the immune system.<br><u>Amino sugars</u>, such as glucosamine and galactosamine, have an amino group substituted for one or more hydroxyl groups in a sugar molecule (...) | *second choice was A. Sugar Acid]* |
| Q16- Which out of the following does not form osazone crystals? | A. Galactose<br>B. Maltose<br>C. Lactose<br>**D. Sucrose** | Sucrose does not form osazone crystals.<br>☐<br>Osazone crystals are formed from the reaction of reducing sugars with phenylhydrazine in the presence of heat. Galactose, maltose, and lactose are reducing sugars and can undergo this reaction to form osazone crystals. However, sucrose is a non-reducing sugar because it does not have a free aldehyde or ketone group, and therefore it cannot form osazone crystals. | Sucrose does not form osazone crystals. ☐ |
| Q17- Which of the following gives a negative reaction to | A. Glucose<br>**B. Maltose**<br>C. Erythrose<br>D. Fructose | Barfoed's test is a test to differentiate between reducing monosaccharides and disaccharides. It involves heating the sugar with Barfoed's reagent, which is a solution of copper acetate in acetic acid. If | **D- Fructose** gives a negative reaction to Barfoed's test ☐ |
| Barfoed's test? |  | the sugar is a reducing sugar, it will react with the copper ions to form a reddish-brown precipitate of copper (I) oxide.<br>Out of the options given, maltose and glucose are reducing sugars, while erythrose and fructose are not. ☐<br>Therefore maltose and glucose would give a positive reaction with Barfoed's test ☐, while erythrose and fructose would give a negative reaction ☐. So the correct answer to the question is option **C**.<br>**Erythrose** ☐ |  |
| Q18- A polysaccharide formed by β1-4 glycosidic linkages between glucose residues is? | A. Inulin<br>B. Amylose<br>C. Agar<br>**D. Cellulose** | The correct answer is D. **Cellulose** ☐<br>Cellulose is a polysaccharide that is composed of repeating units of glucose joined by β1-4 glycosidic linkages (...) | D- **Cellulose** ☐ is a polysaccharide formed by β1-4 glycosidic linkages between glucose residues. Inulin, amylose, and agar do not have β1-4 glycosidic linkages between glucose residues. |
| Q19- Which of the following sugars is | A. Starch<br>**B. Fructose**<br>C. Sucrose | The correct answer is B. **Fructose** ☐ | B- Fructose is predominantly |
| predominantly laevorotatory? | D. Glucose | The term 'laevorotatory' refers to the ability of a compound to rotate the plane of polarised light to the left (counterclockwise) when viewed through a polarimeter (...)<br>Fructose is laevorotatory because it has a chiral centre (an asymmetric carbon atom) that gives it the ability to rotate the plane of polarised light to the left ☐ (...) | laevorotatory ☐ |
| Q20- Which of the following mucopolysaccharides is non-sulfated and most abundant in tissues? | **A. Hyaluronic acid**<br>B. Keratan sulfate<br>C. Heparin<br>D. Dermatan sulfate | The most abundant non-sulfated mucopolysaccharide in tissue is A.<br>**Hyaluronic acid** ☐<br>Hyaluronic acid (also known as hyaluronan) is a linear polysaccharide composed of repeating disaccharides units of N-acetylglucosamine and glucuronic acid. It is a non-sulfated glycosaminoglycan and it is the only glycosaminoglycan that is not covalently attached to proteins to form proteoglycans (...) | The correct answer is A.<br>**Hyaluronic acid.** ☐<br>Hyaluronic acid is a non-sulfated mucopolysaccharide that is highly abundant in tissues, such as in the extracellular matrix and synovial fluid. |

The results in **Table 1** indicate that both ChatGPT and ChatGPT Plus performed well with questions about general knowledge, where abundant and consistent information is likely to be available in the training dataset, for example questions about fundamental chemical properties of the most common mono- and polysaccharides. One peculiar difference is that while the free version of ChatGPT extra text to motivate its MCQ selections, ChatGPT Plus is much less verbose, giving almost always just the selected answer, with no or very little additional text.

For both ChatGPT and ChatGPT Plus we found that one of the key determinants to the bots’ performance is how the question is phrased. ChatGPT and ChatGPT Plus perform generally well with questions asking to describe a feature or property of a sugar and it is seldom able to predict reactivity or classification based on such a description. What neither ChatGPT or ChatGPT Plus can do is to assess if a statement or an answer they give is true or false, which leads both bots astray in questions such as Q2 in **Table 1**: “*Which of the following is a true statement about glucose*”. We found the ChatGPT answer to this question particularly revealing, as it is contradictory and completely nonsensical. Furthermore, as part of the ‘supporting’ statement to its answer to Q2, ChatGPT types “*Glucose has only one asymmetric carbon atom also known as chiral centre*”, which is obviously incorrect, but we thought it may be something worth exploring, as we expected its knowledge of chirality, a basic property of chemical structures, to be better than that. As a note for clarity, the correct answer to Q2 reported in the original resource implies that glucose is in a linear/open chain form. This description often appears in chemistry textbooks, yet the linear form occurs extremely rarely (< 3%) in Nature where glucose is in a cyclic form with five chiral centres.

To investigate this point further we queried about the number of chiral centres in glucose with questions formulated differently within the same thread and also in uncorrelated chats. ChatGPT answered correctly, assuming glucose being in a linear (open chain) conformation, only when the question was phrased as “*How many chiral centres does glucose have?*”. Meanwhile, when we asked “*Does glucose have 4 chiral centres?*”, ChatGPT answers became inconsistent and generally wrong, reporting from 1 to 5 chiral centres in different answers and even listing the different carbon atoms it assumed to be chiral each time. What we believe to be significant to note here is that the structure of the last question is quite similar to the one in Q2, as it asks the bot to answer and to report if its own answer is true or false, which it cannot do, creating a flurry of random and inconsistent statements. The correct and complete answer can be obtained by asking “*How many chiral centres does glucose have in its linear and cyclic forms?*”, which triggers an answer that includes information also on the anomeric centre at C1 generated upon cyclization.

Results from the tests shown in **Table 1** also indicate that ChatGPT/ChatGPT Plus do not perform well in “multi-layered” questions, i.e. questions that require additional, preconceived knowledge to be answered correctly or that touch upon more complex subjects, e.g. topics not extensively covered in resources it was trained on, such as Wikipedia. Q10, Q14 and Q17 are good examples of such questions, where the bots are asked about the expected reactivity of sugars in different laboratory tests. In Q10 ChatGPT claims wrongly that lactose does not form osazone crystals, which it does as it is a reducing sugar, meanwhile in Q14 the correct answer requires knowing what a strong acid is and using this information appropriately to answer the question. The correct answer to Q17 hinges on understanding that maltose is the only disaccharide in the list and that the Barfoed’s test is used to identify monosaccharides. Also interesting how when dealing with less known facts, ChatGPT (but not ChatGPT Plus) fills in random and incorrect information, such as mucic acid being a sugar derived from fructose, yet corrected in a subsequent query (see Q12 in **Table1**), or a dubious description of the natural sources of erythrulose (see Q8 in **Table1**), which is actually a sugar found in red raspberries.

### ChatGPT on writing abstracts

Writing abstracts is a common and frequent task in academia. These short summaries are usually submitted by students and researchers in view of attending scientific meetings, as an advertisement for an invited seminar and/or as part of a published research or review manuscript. The length of an abstract can vary depending on the research field and explicit instructions dictated by publishers or meeting organisers, but generally it does not exceed one A4/letter page (approximately 500 words). For this test we asked ChatGPT to write abstracts on two progressively more specialised topics. Both Topic 1 and 2 explicitly reference published work by one of us[Casalino et al., 2020; Newby et al., 2022], yet while the paper that inspired Topic 1 should be in the ChatGPT training dataset, the paper on which Topic 2 is based has only been published recently. Note, this test was only performed with the free version of ChatGPT. Representative examples of the abstracts we obtained about Topic 1 and 2 are shown in **Table 2**. In terms of formatting, we instructed ChatGPT to limit the abstract to 300 words, which it rarely complied with, to write it in ‘academic style’ and we also explicitly asked for references to complement the text. We intentionally structured the queries to include *verbatim* large portions of the titles of the research articles we sourced the topics from[Casalino et al., 2020; Newby et al., 2022], to check if it would lead to summarising the corresponding papers and/or to plagiarism. As an important note, all the abstracts discussed below were requested in uncorrelated chats and with only slightly varying formats of the queries. The text, content and format of the abstracts obtained in different instances, of which we show representative examples in **Table 2**, seems to be largely the same, while not identical, so fundamentally hinging on the format and wording of the query.

**Table 2.** Two examples of the outputs provided by ChatGPT when asked to write (300 word) abstracts about Topic 1, column 1, and 2, column 2, in academic style. The keywords included in the queries are listed, together with the references given by ChatGPT below the corresponding abstracts. Original (non-prompted) correct and incorrect information is highlighted with green check marks and red crosses, respectively, within the abstract text.

|  |  |
| --- | --- |
| **Topic 1.** *“Write an abstract of 300 words in academic style with references about the role of the glycan shield in the SARS-CoV-2 spike protein”* | **Topic 2.** *“Write an abstract of 300 words in academic style with references about how variations within the glycan shield of SARS-CoV-2 impact the spike dynamics”* |
| **Keywords in query:** glycan shield, SARS-CoV-2, spike protein | **Keywords in query:** variations, glycan shield, SARS-CoV-2, spike, spike dynamics |
| **ChatGPT Abstract.** The glycan shield, composed of complex carbohydrate structures known as glycans, plays a crucial role in the biology of the severe acute respiratory syndrome coronavirus 2 (SARS-CoV-2), the virus responsible for the ongoing COVID-19 pandemic. The spike (S) protein on the surface of SARS-CoV-2, which facilitates viral entry into host cells, is heavily glycosylated with glycans that decorate its surface □. These glycans act as a shield, concealing vulnerable epitopes and protecting the virus from immune recognition, thus facilitating viral immune evasion and increasing its infectivity□. In this review □, we highlight the diverse roles of glycans in the SARS-CoV- | **ChatGPT Abstract.** SARS-CoV-2, the virus responsible for the COVID-19 pandemic, has a complex glycan shield that plays a significant role in the virus’s ability to evade the host immune system. Variations within the glycan shield can impact the dynamics of the viral spike, which is the protein that allows the virus to enter the host cell. Understanding how variations of the glycan shield impact viral spike dynamics is crucial for the development of effective antiviral strategies.<br>Studies have shown that the presence of certain glycans on the viral spike can alter its stability, conformation and interaction with the host cells □. For example the presence of sialylated glycan on the viral spike has been found to increase the stability of the spike □ and enhance the virus ability to enter host |
| 2 S protein and their implications in viral pathogenesis.<br>First, we discuss the structural diversity and abundance of glycans on the SARS-CoV-2 S protein. The glycosylation sites on the S protein are numerous and strategically located, particularly in the receptor-binding domain (RBD) □, which is responsible for binding to the host cell receptor, angiotensin-converting enzyme 2 (ACE2). The glycans on the S protein are predominantly of the complex type, with extensive branching and sialylation □, making them highly adaptable and capable of shielding the virus from immune surveillance.<br>Next, we delve into the functional roles of the glycan shield in the SARS-CoV-2 S protein. The glycans act as a physical barrier, preventing direct contact between the viral protein and host immune cells, thereby reducing antibody recognition and neutralization. Additionally, the glycans on the S protein can modulate viral infectivity by influencing the conformational dynamics and stability of the RBD □, affecting ACE2 binding and viral fusion.<br>Moreover, the glycans can also engage in molecular mimicry, resembling host glycans and evading immune detection □. | cells□. On the other hand the presence of fucosylated glycans has been associated with reduced stability and reduced infectivity of the virus □.<br>Variation of the glycan shield of SARS-CoV-2 can also impact the ability of neutralizing antibodies to bind to the viral spike □. Antibodies targeting the viral spike play a critical role in the host immune response to SARS-CoV-2. However, the presence of certain glycans on the viral spike can prevent the binding of neutralizing antibodies, leading to increased viral persistence and infectivity.□<br>Furthermore, variations in the glycan shield can impact the ability of the virus to evade the immune system by changing the molecular recognition of the virus by the host immune system. This can result in changes to the virus's susceptibility to immune-mediated clearance and its ability to establish persistent infections.<br>In conclusion, variations within the glycan shield of SARS-CoV-2 have a significant impact on viral spike dynamics, including the stability and conformation of the spike, its interactions with host cells, and its |
| Furthermore, we discuss the implications of the glycan shield in the development of COVID-19 therapeutics and vaccines. The glycosylation patterns on the S protein are subject to variation, and mutations in the viral genome can alter glycan occupancy and composition, affecting viral infectivity and immune recognition □. Understanding the role of glycans in the SARS-CoV-2 S protein is critical for the design and optimization of antiviral strategies, including vaccines and therapeutics that target the glycan shield to enhance immune response and neutralization of the virus.<br>In conclusion, the glycan shield on the SARS-CoV-2 S protein plays multifaceted roles in viral immune evasion, infectivity, and pathogenesis. Further research on the composition, structure, and function of glycans on the S protein is warranted to elucidate their precise mechanisms and exploit them for the development of effective COVID-19 interventions. **[401 words]** | susceptibility to immune mediated clearance.<br>Understanding these interactions is crucial for the development of effective antiviral strategies to combat SARS-CoV-2 and other related viruses. □ **[321 words]** |
| **ChatGPT List of References.**<br>1. Watanabe Y, Allen JD, Wrapp D, McLellan JS, Crispin M. Site-specific glycan analysis of the SARS-CoV-2 spike. Science. 2020 □ Sep 11;369(6500):330-333. □ | **ChatGPT List of References.**<br>1. E. A. Hernandez, J.J. van Dongen, and M.A. Peppelenbosh, "The Dynamic Glycan Shield of SARS-CoV-2: Implication for Virus Biology and Immune Evasion" Trends in Microbiology vol. 29, no. |
| 2. Casalino L, Gaieb Z, Goldsmith JA, Hjorth CK, Dommer AC, Harbison AM, Fogarty CA, Barros EP, Taylor BC, McLellan JS, Fadda E, Amaro RE. Beyond shielding: the roles of glycans in the SARS-CoV-2 spike protein. ACS Cent Sci. □ 2021 Mar 24;7(3):471-479. □<br>3. Vankadari N, Wilce JA. Emerging WuHan (COVID-19) coronavirus: glycan shield and structure prediction of spike glycoprotein and its interaction with human CD26. Emerg Microbes Infect. 2020 □ Feb 21 □;9(1):601-604 □ | 7, pp 589-598 (2021) □<br>2. K.M. Vermaas, B.W. Kroeze, M.A. Peppelenbosh, and J.J. van Dongen "SARS-CoV-2 Spike Glycoprotein: The Role of N-Glycans in Viral Biology and Antiviral Strategies" Viruses, vol. 13, no. 3, pp 321 (2021) □ |

In terms of original content, the ChatGPT abstract on Topic 1 is marginally of higher quality than the one on Topic 2, which could be due to the fact that the paper that inspired Topic 1 may be included in the training dataset[Casalino et al., 2020], or to the broader breadth of the subject. Indeed, ChatGPT defined correctly what the glycan shield is, what is the spike protein, and mentions the role of the glycan shield in immune evasion, molecular mimicry and in increasing viral infectivity. There is also a mention of its role in affecting the spike’s conformational dynamics and stability, although no explanation on how that is so. However, this potentially useful content is presented in an disjointed and repetitive format, with an unwarranted indication that the abstract is for a “review”. The abstract also includes parts that can be defined as “hallucinations”, reporting on hyper-sialylation of the spike and high branching levels of complex N-glycans responsible for more effective shielding. In regard to the references listed, all correspond to existing work, and 1 and 2 are almost correct, except for the page, volume and issue numbers. Reference 3 appears to be correct, except for the inclusion of “Feb”, which possibly matches the article’s final submission date, rather than the publication date.

As an interesting detail, when we included “with references” in the queries, as shown in the examples in **Table 2**, references were listed, sometimes formatted differently and with varying degrees of correctness, see also the subsection on essays below, however they were never cited within the text. The specification of a precise reference format in the query, such as “with Harvard references”, led to adding the citations in the text. We also asked to include the corresponding Digital Object Identifiers (DOIs) to the references and in every test we performed the listed DOI is either made-up (most frequently) or corresponds to a completely unrelated paper. So far we have not obtained a DOI from ChatGPT that corresponds to the paper in the same reference, unless we gave DOIs ourselves in the query.

In terms of content, the overall quality of the abstract on Topic 2 is rather poor, which may be due to the higher level of specificity of the subject, relative to Topic 1, and/or the fact that the work that inspired the query was published only recently[Newby et al., 2022], and thus it is not included in the bot’s training dataset. Indeed, ChatGPT is not able to assign the appropriate meaning to “variations within the glycan shield”. The correct information provided in this abstract relates to the ability of the glycan shield to alter the spike’s stability and dynamics and to hinder its recognition by the immune system. Yet, as in the other abstract, this information is presented in a non-cohesive, vague and repetitive framework. Also, it is embedded with fabricated content stating that while sialylation increases stability of the spike and the infectivity of the virus, fucosylation decreases stability and infectivity. Furthermore, the references listed in this abstract are also completely made-up, from the list of authors, to the titles, to the publication dates and journal details. Yet, the names of the journals correspond to known scientific publications, which may give a false sense of security to the distracted reader.

Finally, in additional tests we asked ChatGPT to write an abstract (with the same or similar formatting conditions described above) or to summarise (with no formatting conditions) the two published papers that inspired Topic 1[Casalino et al., 2020] and 2[Newby et al., 2022], among others. None of these tests led to obvious (or detectable with absolute certainty) levels of plagiarism, but did not lead to accurate descriptions of those papers either and still included unwarranted fabrications, such as the claim that the B.1.1.7 variant (alpha) carried extensive mutations of the S glycan shield, or that the reference[Newby et al., 2022] described experiments that in fact were never part of the work.

### ChatGPT on writing essays

With this test we wanted to explore if and how ChatGPT could address the writing of longer and more complex documents, such as essays, often an integral part of take-home assignments in higher education. Also in this case all tests were performed only with the free version of ChatGPT. As a *caveat*, ChatGPT is not built to write essays, having an unwritten (unofficial) 4000 characters limit, corresponding to approximately 500 words. Our attempts at making the bot write longer text by enforcing a “no less than 2000 words” limit in the query have not been successful, allowing it to reach from 550 to 750 words maximum. Note: when the word limit is reached, the user can instruct ChatGPT to continue from where it stopped, which it does seamlessly. Also, the text-length limitation does not preclude the user to compile an essay from multiple separate sections, which we did not attempt in this context as we believed to be outside the scope of this work. Instead, we asked ChatGPT to write (short) essays in the form of literature reviews.

The most complex topics we tested ChatGPT on are, Essay 1 titled “*The role of glycosylation in enveloped viruses infection*” and Essay 2 titled “*Automated synthesis of complex glycans*”. For each of these, we specified a list of key references to be included in the text. Representative versions of the two essays and corresponding references are included as Supplementary Material. Both outputs are far from satisfactory in terms of content and structure, regardless of their short length. The output of Essay 1 can be described as a combined version of the abstracts analysed in the previous subsection, where the broad lines of the paper ChatGPT should have in its training dataset[Casalino et al., 2020] are reported, again without any level of in-depth knowledge, understanding or critical analysis. Meanwhile, the description of the second paper[Newby et al., 2022], published recently and therefore not in the training dataset, is even more vague and filled with inaccuracies and fabrications. Essay 2 is slightly longer than Essay 1, yet the content is as generic, highly hinging on the titles of the papers to produce text and dotted with inaccuracies. For example, in the (short) description of the 2003 review by P. Seeberger on automated glycan synthesis[Seeberger, 2003], it states “*The author described the development of glycan microarrays*”, which is not part of that review in any form.

Examples of outputs we obtained from more descriptive queries, focusing on general and heavily documented subjects in Glycoscience, such as the biosynthetic pathways leading to N- and O-glycosylation of proteins, are also shown as Supplementary Material. These range from vague and superficial to factually wrong, e.g. in Essay 3 “… *assembly of the oligosaccharide on the LLO precursor. This process is initiated by the transfer of a GlcNAc residue from UDP-GlcNAc to the dolichol-linked Man5GlcNAc2-PP-dolichol by the oligosaccharyltransferase (OST) complex*”. The test is complemented by completely fabricated references that surprisingly include the names of highly recognizable leaders in the field, with plausible titles, and names of real journals, which may give a false aura of credibility to a distracted reader.

## Discussion

The series of tests we presented and analysed in the previous section are limited to specific fields of Science, namely Carbohydrate Chemistry and Glycobiology, and also cover a potentially limited set of academic writing formats, i.e. short exam questions, MCQs and short (less than 1000 words) abstracts/summary pieces. Yet, we believe that the results we obtained may shed some light on the potentials and limitations of the currently available free OA version of ChatGPT (based on GPT-3.5) in this context, with points of reflection easily applicable to other areas of Chemistry and Life Sciences.

In terms of potentials, ChatGPT did generally well within the boundaries of what it was built and trained to do, namely answering descriptive questions, in a fashion that is pleasantly discursive, while being able to predict outcomes from such descriptions. Because of its extensive training, which is likely to include most basic chemistry and biology, supplemented by plenty of correct information on those subjects, it scored very high (70%) on a basic Carbohydrate Chemistry MCQ test where the majority of questions were descriptive. As a point of comparison we have run the same MCQ deck with ChatGPT Plus, a new-and-improved subscription only service that became available only recently. The performance was surprisingly worse, with only 55% of correct responses. Also, ChatGPT can write short and generally accurate descriptors about subjects that are part of sufficiently broad knowledge, i.e. extensively discussed in the media, books, Wikipedia or other web resources it is trained on, and it can do that in virtually any writing format or style. In this work we presented the results of tests using exclusively an academic style, but we also asked ChatGPT to rewrite text as a hip-hop or rap song and in Shakespearean English (not included here), with interesting potential applications in science communication and public outreach.

Science though is a tricky subject for this type of applications of LLMs, because most scientific writing does not involve just broad descriptions of phenomena, but most often it requires critical assessment based on data analysis. Because it is an LLM, ChatGPT has no ability to “think”; it does not know whether the information it gives is true or false, as its outputs are sequences of words built on probabilities, with weights dependent on training sets and algorithm design. As N. Chomsky *et al*. eloquently stated in a recent opinion piece in the New York Times[Chomsky et al.] “*The correct explanations of language are complicated and cannot be learned just by marinating in big data*.” Therefore, as we have seen from our potentially limited tests, when ChatGPT is asked questions in a format that requires a “true or false” assessment of its own output, or implies previous knowledge, or highly specialised knowledge, or information that is not explicitly part of its training set, it generally fails. The dangerous limitation of the model is that it cannot alert the user when the output is fabricated, as it has no idea if and when that may be. As we have seen in our tests, these fabrications, or “hallucinations” are virtually everywhere, from dotting largely correct outputs, to sidetracking the whole narrative of others.

Based on these considerations, we strongly believe that all ChatGPT users should be aware of these shortcomings and use the tool with extreme caution when studying, researching topics or drafting text. We believe that the use of ChatGPT may be particularly treacherous for students using it in preparation for exams. In fact, while written exams are generally held in presence in halls or classrooms where students work alone, allowed only to use pen and paper, monitored by invigilators, ChatGPT could be seen as a shortcut to prepare for exams, rather than studying from handouts, notes or books, by querying previous exam questions and memorising answers.

From the educators point of view, it is highly unlikely that any bot built to detect plagiarism or wording probabilities will be able to flag with a high degree of certainty a text produced by AI. So, it seems to us that it will be useful to learn how to work with ChatGPT, using it to our advantage. As an example, from an assessment point-of view we can think of LLMs as tools to formulate exam questions in different formats. Our tests show that ChatGPT and ChatGPT Plus are very efficient at that when they have a good grasp on the topic. For example ChatGPT Plus (chosen here because its answers are less verbose) can easily create an MCQ on a topic it knows, such as “what is the most abundant sugar in honey” see Q6 in **Table 1**, with one correct answer and as many decoy options as needed. This strategy can be exported to any topic and any question format, exploiting ChatGPT’s keen ability to adapt text to different formats and styles.

From the teaching and learning point of view, with this project we demonstrated an example inspired by a “flipped class” model. Here the student (DOW), under the lecturer’s supervision (EF), was asked to assess the scientific content produced by ChatGPT, to determine if the information the bot provided was true or false through independent studying, and to try to devise strategies to obtain complete or correct information wherever possible. Furthermore, through this effort, we also learned that the much feared negative impact of ChatGPT in assessments can be curbed by rephrasing questions in a way that the bot cannot answer, by always asking for a critical assessment, rather than descriptors and predictions. As a final note, easily applicable in the field of Chemistry, exam questions that involve drawing chemical structures, reactions schemes and/or pathways, completely bypass the current version of ChatGPT, as it can only produce text, and any ability of currently available AI drawing tools, such as DALL-E and DALL-E-2 (https://openai.com), which are built for different purposes.

## Conclusions

In this work we presented and discussed the performance of the LLM ChatGPT in addressing progressively more complex and specialised questions in Carbohydrate Chemistry and Glycobiology. Based on the results we obtained, we found that ChatGPT can generally answer correctly short and descriptive questions about general and basic knowledge, as those are likely to be heavily documented in its training dataset. In some cases it can also elaborate simple predictions and infer consequences based on the descriptors it is given or it is trained on, yet it cannot evaluate or make assessments.

We also found that in virtually all the tests we performed, the output was likely to contain fabricated content. While answering short and descriptive queries the made-up content appeared sporadically, yet it represented the majority of the answer when dealing with complex, non-descriptive and highly specialised subjects. Here we propose that the knowledge of these shortcomings can be used to guide the phrasing of exams questions and the structuring of writing-based assessments in higher education. We also believe that this work may represent a useful example of how ChatGPT can be used within a “flipped class” model, hopefully inspiring further exploration of the integration of LLMs in teaching and learning in higher education.

## Notes and Acknowledgements

DOW is a final year BSc Double Honors student in Chemistry and in Mathematics and Statistics at Maynooth University. The work presented in this manuscript is part of DOW’s final year project in Chemistry. EF is an Associate Professor in the Department of Chemistry, affiliated to the Hamilton Institute at Maynooth University and project supervisor. EF designed the project. DOW ran all the tests on ChatGPT and reported the results. DOW and EF analysed, discussed and summarised the results. DOW and EF wrote the manuscript. DOW and EF gratefully acknowledge Prof David Malone from the Department of Mathematics and Statistics and Hamilton Institute at Maynooth University for insightful discussions.

## Notes

### Competing Interest Statement

The authors have declared no competing interest.

### Summary of Updates

Table 1 Includes results obtained by testing the performance of both, the free version of ChatGPT and the subscription-only version ChatGPT Plus for comparison, on a set of 20 MCQs in Carbohydrate Chemistry

## References

Bommasani R, Hudson DA, Adeli E, Altman R, Arora S, von Arx S, Bernstein MS, Bohg J, Bosselut A, Brunskill E, Brynjolfsson E, Buch S, Card D, Castellon R, Chatterji N, Chen A, Creel K, Davis JQ, Demszky D, Donahue C, Doumbouya M, Durmus E, Ermon S, Etchemendy J, Ethayarajh K, Fei-Fei L, Finn C, Gale T, Gillespie L, Goel K, Goodman N, Grossman S, Guha N, Hashimoto T, Henderson P, Hewitt J, Ho DE, Hong J, Hsu K, Huang J, Icard T, Jain S, Jurafsky D, Kalluri P, Karamcheti S, Keeling G, Khani F, Khattab O, Koh PW, Krass M, Krishna R, Kuditipudi R, Kumar A, Ladhak F, Lee M, Lee T, Leskovec J, Levent I, Li XL, Li X, Ma T, Malik A, Manning CD, Mirchandani S, Mitchell E, Munyikwa Z, Nair S, Narayan A, Narayanan D, Newman B, Nie A, Niebles JC, Nilforoshan H, Nyarko J, Ogut G, Orr L, Papadimitriou I, Park JS, Piech C, Portelance E, Potts C, Raghunathan A, Reich R, Ren H, Rong F, Roohani Y, Ruiz C, Ryan J, Ré C, Sadigh D, Sagawa S, Santhanam K, Shih A, Srinivasan K, Tamkin A, Taori R, Thomas AW, Tramèr F, et al. 2021. On the Opportunities and Risks of Foundation Models. arXiv [cs.LG].

Casalino L, Gaieb Z, Goldsmith JA, Hjorth CK, Dommer AC, Harbison AM, Fogarty CA, Barros EP, Taylor BC, McLellan JS, Fadda E, Amaro RE. 2020. Beyond Shielding: The Roles of Glycans in the SARS-CoV-2 Spike Protein. ACS Cent Sci 6: 1722–1734.

Chomsky N, Roberts I, Watumull J, Chomsky N. The false promise of ChatGPT. NY Times.

Ferruz N, Schmidt S, Höcker B. 2022. ProtGPT2 is a deep unsupervised language model for protein design. Nat. Commun. 13: 4348.

Lin Z, Akin H, Rao R, Hie B, Zhu Z, Lu W, Smetanin N, Verkuil R, Kabeli O, Shmueli Y, Dos Santos Costa A, Fazel-Zarandi M, Sercu T, Candido S, Rives A. 2023. Evolutionary-scale prediction of atomic-level protein structure with a language model. Science 379: 1123–1130.

Madani A, Krause B, Greene ER, Subramanian S, Mohr BP, Holton JM, Olmos JL Jr, Xiong C, Sun ZZ, Socher R, Fraser JS, Naik N. 2023. Large language models generate functional protein sequences across diverse families. Nat. Biotechnol.

Newby ML, Fogarty CA, Allen JD, Butler J, Fadda E, Crispin M. 2022. Variations within the Glycan Shield of SARS-CoV-2 Impact Viral Spike Dynamics. J. Mol. Biol. 435: 167928.

Seeberger PH. 2003. Automated carbohydrate synthesis to drive chemical glycomics. Chem. Commun.: 1115–1121.

Stokel-Walker C, Van Noorden R. 2023. What ChatGPT and generative AI mean for science. Nature 614: 214–216.

Vaswani A, Shazeer N, Parmar N, Uszkoreit J, Jones L, Gomez AN, Kaiser Ł, Polosukhin I. 2017. Attention is all you need. Adv. Neural Inf. Process. Syst. 30.

Vu MH, Akbar R, Robert PA, Swiatczak B, Sandve GK, Greiff V, Haug DTT. 2023. Linguistically inspired roadmap for building biologically reliable protein language models. Nature Machine Intelligence 5: 485–496.

