## Supplementary Material for "Can ChatGPT pass Glycobiology?"

**ChatGPT on writing essays**. Here below we show representative examples of the outputs that ChatGPT gave when we asked to write literature reviews, as a format model for essays.

For this task we chose the following titles, 1 “*The importance of glycosylation in enveloped viruses*” and 2 “*Automated synthesis of complex glycans*”, as examples of challenging writing pieces. Note, in these in all these tests we did not specify a limit for the text’s length because, based on our reading, it appears that ChatGPT has “a hidden” limit of 500 words or of approximately 4000 characters. For this reason, when we enforce in the query to keep a word count exceeding this limit, e.g. stating “no less than 2000 words”, ChatGPT truncates the text mid-sentence after approximately 500 words. As shown in Essay 2 below, when the user asks in the chat to continue the section after the break, the text is usually resumed seemslessly.

Furher below we show examples (Essay 3 and 4) of outputs produced from queries requesting descriptions of topics in basic Glycobiology, such as “*Describe the biosynthetic pathway leading to N-glycosylation of proteins*” and “*Describe the biosynthetic pathway leading to O-glycosylation of proteins*”. Despite the easier subject and the descriptive nature of the query the outputs also contain serious mistakes and inaccuracies and are complemented by completely fabricated references.

**Essay 1.** *The importance of glycosylation in enveloped viruses infection*

**Query.** *Write a literature review in an academic style including Harvard references with the title the importance of glycosylation in enveloped viruses infection. Including the following references,*

*Lorenzo Casalino, Zied Gaieb, Jory A. Goldsmith, Christy K. Hjorth, Abigail C. Dommer, Aoife M. Harbison, Carl A. Fogarty, Emilia P. Barros, Bryn C. Taylor, Jason S. McLellan, Elisa Fadda, and Rommie E. Amaro, Beyond Shielding: The Roles of Glycans in the SARS-CoV-2 Spike Protein, ACS Central Science 2020 6 (10), 1722-1734 DOI: 10.1021/acscentsci.0c01056*

*Newby ML, Fogarty CA, Allen JD, Butler J, Fadda E, Crispin M. Variations within the Glycan Shield of SARS-CoV-2 Impact Viral Spike Dynamics. J Mol Biol. 2023 Feb 28;435(4):167928. DOI: 10.1016/j.jmb.2022.167928*

**ChatGPT Output.**

Title: The Importance of Glycosylation in Enveloped Viruses

Glycosylation, the process of adding sugar molecules to proteins or lipids, plays a critical role in the structure and function of enveloped viruses. Enveloped viruses, such as SARS-CoV-2, the virus responsible for the COVID-19 pandemic, are surrounded by a lipid bilayer derived from the host cell membrane, which is studded with viral glycoproteins. These glycoproteins are often heavily glycosylated, meaning they are decorated with complex sugar moieties. The glycan shield created by these glycans on viral surface proteins is thought to be involved in various aspects of viral infection, including immune evasion, viral entry, and viral protein stability.

One of the key findings regarding the importance of glycosylation in enveloped viruses comes from recent research on SARS-CoV-2, the virus responsible for the COVID-19 pandemic. Casalino et al. (2020) used molecular dynamics simulations and computational modeling to study the role of glycans in the spike protein of SARS-CoV-2, which is responsible for viral entry into host cells. They found that the glycans on the spike protein play a crucial role in stabilizing the protein structure, shielding it from immune recognition, and modulating the conformational dynamics of the spike protein. The authors suggest that the glycans on the spike protein of SARS-CoV-2 are not just a passive shield, but rather active participants in viral infectivity.

Furthermore, Newby et al. (2023) conducted a study investigating the impact of glycan variations on the dynamics of the spike protein of SARS-CoV-2. Using molecular simulations and bioinformatics analysis, they showed that even small changes in the glycan shield of the spike protein can have significant effects on its stability and dynamics, potentially impacting viral infectivity. This study highlights the dynamic nature of viral glycans and their contribution to viral protein function.

In addition to SARS-CoV-2, glycosylation has been shown to be important in other enveloped viruses as well. For example, in HIV, the viral envelope glycoprotein gp120 is heavily glycosylated and the glycans on gp120 have been implicated in viral entry, immune evasion, and resistance to neutralizing antibodies. In influenza viruses, the hemagglutinin protein is also heavily glycosylated and the glycans on hemagglutinin play a role in viral entry, antigenicity, and immune evasion (Newby et al., 2023).

In conclusion, glycosylation plays a crucial role in enveloped viruses, including SARS-CoV-2, HIV, and influenza viruses, by contributing to viral protein stability, immune evasion, and viral entry. Further research in this area is needed to better understand the precise mechanisms by which glycans modulate viral infectivity and to potentially develop novel antiviral strategies targeting viral glycosylation. Future studies could also explore the interplay between viral glycans and host glycans, as well as the impact of glycosylation on other aspects of viral replication, such as viral assembly and release. **[449 words]**

References:

1. Lorenzo Casalino, Zied Gaieb, Jory A. Goldsmith, Christy K. Hjorth, Abigail C. Dommer, Aoife M. Harbison, Carl A. Fogarty, Emilia P. Barros, Bryn C. Taylor, Jason S. McLellan, Elisa Fadda, and Rommie E. Amaro. Beyond Shielding: The Roles of Glycans in the SARS-CoV-2 Spike Protein. ACS Central Science 2020, 6 (10), 1722-1734. DOI: 10.1021/acscentsci.0c01056.
2. Newby ML, Fogarty CA, Allen JD, Butler J, Fadda E, Crispin M. Variations within the Glycan Shield of SARS-CoV-2 Impact Viral Spike Dynamics. J Mol Biol. 2023 Feb 28;435(4):167928. doi: 10.1016/j.jmb.2022.167928.

**Essay 2.** *Automated synthesis of complex glycans*

**Query.** *Write a literature review about the automated synthesis of complex glycans using all four references below:*

*Yao, W., Xiong, DC., Yang, Y. et al. Automated solution-phase multiplicative synthesis of complex glycans up to a 1,080-mer. Nat. Synth 1, 854–863 (2022).* [*https://doi.org/10.1038/s44160-022-00171-9*](https://doi.org/10.1038/s44160-022-00171-9) *,*

*Wu, Y., Xiong, DC., Chen, SC. et al. Total synthesis of mycobacterial arabinogalactan containing 92 monosaccharide units. Nat Commun 8, 14851 (2017).* [*https://doi.org/10.1038/ncomms14851,*](https://doi.org/10.1038/ncomms14851,)

*Panza M, Pistorio SG, Stine KJ, Demchenko AV. Automated Chemical Oligosaccharide Synthesis: Novel Approach to Traditional Challenges. Chem Rev. 2018 Sep 12;118(17):8105-8150.* [*https://doi.org/10.1021/acs.chemrev.8b00051*](https://doi.org/10.1021/acs.chemrev.8b00051)

*Seeberger PH. Automated carbohydrate synthesis to drive chemical glycomics. Chem Commun (Camb). 2003 May 21;(10):1115-21.* [*https://doi.org/10.1039/b210230g*](https://doi.org/10.1039/b210230g)

*In the review include an introduction, main body, conclusions and Harvard references.*

**ChatGPT Output**

Title: Automated Synthesis of Complex Glycans: Advancements and Applications

Introduction: Glycans, also known as carbohydrates or sugars, are essential biomolecules that play crucial roles in various biological processes, including cell-cell recognition, signaling, and immune responses. The structural complexity and diversity of glycans make their chemical synthesis challenging, requiring precise control over stereochemistry, protecting group strategies, and glycosylation reactions. Traditional methods of glycan synthesis are labor-intensive, time-consuming, and often result in low yields, limiting the availability of complex glycans for functional studies and therapeutic applications.

In recent years, there has been significant progress in the field of automated synthesis of complex glycans, overcoming many of the limitations of traditional methods. Automated synthesis platforms provide precise control over reaction conditions, reduce human error, and enable the synthesis of larger and more complex glycans with higher yields. In this literature review, we will discuss recent advancements in the field of automated synthesis of complex glycans, focusing on the four key references: Yao et al. (2022), Wu et al. (2017), Panza et al. (2018), and Seeberger (2003).

Main Body: Yao et al. (2022) reported an innovative approach for automated solution-phase multiplicative synthesis of complex glycans up to a 1,080-mer. Using a glycan synthesis robot, the authors achieved high yields and excellent stereoselectivity in the synthesis of a variety of complex glycans, including branched and linear structures. The automated synthesis platform allowed for precise control over reaction conditions, enabling the synthesis of glycans with diverse structures and functionalities. This approach opens up new possibilities for the synthesis of large and complex glycans, which were previously challenging to access using traditional methods.

Wu et al. (2017) described a total synthesis of mycobacterial arabinogalactan, a complex glycan containing 92 monosaccharide units, using an automated solid-phase synthesis approach. The authors utilized a strategic combination of protecting group strategies and glycosylation reactions to achieve the synthesis of this large and structurally diverse glycan. The automated solid-phase synthesis platform enabled efficient and scalable synthesis of this complex glycan, providing a valuable tool for studying the structure and function of arabinogalactan and its role in mycobacterial cell wall assembly.

Panza et al. (2018) reviewed the advancements in automated chemical oligosaccharide synthesis, providing a comprehensive overview of the strategies, challenges, and applications of this approach. The authors discussed the use of solid-phase and solution-phase automated synthesis platforms for the synthesis of diverse types of glycans, including linear and branched structures, sialylated glycans, and glycosaminoglycans. The review highlighted the advantages of automated synthesis, such as improved precision, scalability, and reproducibility, and its potential for driving chemical glycomics research.

Seeberger (2003) discussed the application of automated carbohydrate synthesis in the field of chemical glycomics. The author described the development of glycan microarrays, which enable high-throughput screening of glycan-protein interactions, and the use of automated synthesis platforms for the preparation of diverse glycan libraries for screening purposes. The review highlighted the importance of automated synthesis in advancing our understanding of glycan function and its potential for accelerating drug discovery efforts targeting glycan-protein interactions.

Conclusions: The automated synthesis of complex glycans has emerged as a powerful tool in glycoscience, enabling the synthesis of diverse and structurally complex glycans with higher yields, improved precision, and scalability. The advancements in automated synthesis platforms have overcome many of the limitations of traditional glycan synthesis methods and opened up new possibilities for the study of glycan structure and (….)^[2]^ function, as well as for the development of glycan-based therapeutics and diagnostics. The references Yao et al. (2022), Wu et al. (2017), Panza et al. (2018), and Seeberger (2003) collectively highlight the significant progress made in the field of automated synthesis of complex glycans, ranging from solution-phase multiplicative synthesis to solid-phase synthesis, and their diverse applications in glycoscience research, chemical glycomics, and drug discovery.

In conclusion, automated synthesis of complex glycans has revolutionized the field of glycoscience, providing efficient and scalable methods for the synthesis of diverse glycans with high precision and reproducibility. These advancements have facilitated the access to complex glycans, which were previously challenging to synthesize using traditional methods, and have expanded our understanding of glycan structure and function. Automated synthesis platforms have also enabled the development of glycan-based therapeutics and diagnostics, as well as high-throughput screening of glycan-protein interactions. Overall, automated synthesis of complex glycans has the potential to drive further advancements in glycoscience research and accelerate the discovery of novel glycan-based therapies. **[715 words]**

Harvard References:

1. Yao, W., Xiong, DC., Yang, Y. et al. Automated solution-phase multiplicative synthesis of complex glycans up to a 1,080-mer. Nat. Synth 1, 854–863 (2022). <https://doi.org/10.1038/s44160-022-00171-9>
2. Wu, Y., Xiong, DC., Chen, SC. et al. Total synthesis of mycobacterial arabinogalactan containing 92 monosaccharide units. Nat Commun 8, 14851 (2017). <https://doi.org/10.1038/ncomms14851>
3. Panza M, Pistorio SG, Stine KJ, Demchenko AV. Automated Chemical Oligosaccharide Synthesis: Novel Approach to Traditional Challenges. Chem Rev. 2018 Sep 12;118(17):8105-8150. <https://doi.org/10.1021/acs.chemrev.8b00051>
4. Seeberger PH. Automated carbohydrate synthesis to drive chemical glycomics. Chem Commun (Camb). 2003 May 21;(10):1115-21. <https://doi.org/10.1039/b210230g>

**Essay 3**. *Describe the biosynthetic pathway leading to the N-glycosylation of proteins*

**Query.** *Describe the biosynthetic pathway leading to the N-glycosylation of proteins that include Harvard references with corresponding DOIs.*

**ChatGPT Output**

N-glycosylation is a post-translational modification that involves the addition of a glycan chain to an asparagine residue in a protein. The biosynthetic pathway leading to N-glycosylation is a complex process that involves the sequential action of several enzymes. Here, we describe the biosynthetic pathway leading to N-glycosylation in proteins based on recent literature.

1 Synthesis of the sugar precursor: The first step in N-glycosylation is the synthesis of the lipid-linked oligosaccharide (LLO) precursor. This process takes place in the endoplasmic reticulum (ER) membrane and involves the sequential addition of monosaccharides to a dolichol phosphate carrier. The LLO precursor is then flipped to the luminal side of the ER membrane for further processing (1).

2 Assembly of the oligosaccharide: The second step in N-glycosylation involves the assembly of the oligosaccharide on the LLO precursor. This process is initiated by the transfer of a GlcNAc residue from UDP-GlcNAc to the dolichol-linked Man5GlcNAc2-PP-dolichol by the oligosaccharyltransferase (OST) complex. This is followed by the sequential addition of several mannose and glucose residues, which is mediated by specific glycosyltransferases. The final oligosaccharide structure, Man9GlcNAc2, is recognized by the lectin chaperones calnexin and calreticulin, which assist in the proper folding of the nascent protein (2).

3 Transfer of the oligosaccharide to the protein: The third step in N-glycosylation involves the transfer of the oligosaccharide from the LLO precursor to the protein. This is catalyzed by the OST complex, which transfers the oligosaccharide to an asparagine residue within the consensus sequence Asn-X-Ser/Thr (where X is any amino acid except proline). The transfer occurs cotranslationally as the nascent polypeptide chain is being translocated into the ER lumen by the translocon (3).

In summary, the biosynthetic pathway leading to N-glycosylation in proteins involves the synthesis of a lipid-linked oligosaccharide precursor, assembly of the oligosaccharide on the LLO precursor, and transfer of the oligosaccharide to the protein by the OST complex. The precise mechanisms and enzymes involved in each step of the pathway have been the subject of extensive research over the years and continue to be an active area of investigation.

References:

1 Aebi, M. et al. N-Linked Protein Glycosylation in the ER. Biochemistry DOI: 10.1021/acs.biochem.6b00707 (2017).

2 Kelleher, D. J. & Gilmore, R. An evolving view of the eukaryotic oligosaccharyltransferase. Glycobiology DOI: 10.1093/glycob/cwaa030 (2020).

3 Pfeffer, S. & Aebi, M. Intracellular Glycosylation of Proteins. Glycobiology DOI: 10.1093/glycob/cwaa054 (2020).

**Notes of the Authors:** There are glaring factual mistakes in the text produced, such as disjoining steps 1 and 2, which in fact are the same step, stating that “*This process is initiated by the transfer of a GlcNAc residue from UDP-GlcNAc to the dolichol-linked Man5GlcNAc2-PP-dolichol by the oligosaccharyltransferase (OST) complex*”, which in fact is a step that does not occur, the omission of the nature/sequence of the actual dolichol-linked precursor transferred to the protein by OST. Also N-glycosylation is a posttranslational modification that can occur cotranslationally.

Furthermore, despite the references listed cite name of well known authors, leaders in this specific area of research, these are entirely fabricated, from the titles and dates of publication, to the DOIs, which in this specific example correspond all to different articles. The names of the journals correspond to existing journals.

**Essay 4**. *Describe the biosynthetic pathway leading to the O-glycosylation of proteins*

**Query.** *Describe the biosynthetic pathway leading to the O-glycosylation of proteins that include Harvard references with corresponding DOIs.*

**ChatGPT Output**

O-glycosylation is a post-translational modification that involves the addition of a glycan chain to a serine or threonine residue in a protein. The biosynthetic pathway leading to O-glycosylation is a complex process that involves the sequential action of several enzymes. Here, we describe the biosynthetic pathway leading to O-glycosylation in proteins based on recent literature.

1 Initiation of O-glycosylation: The first step in O-glycosylation involves the addition of a single N-acetylgalactosamine (GalNAc) residue to a serine or threonine residue in a protein. This reaction is catalyzed by a family of enzymes known as UDP-GalNAc:polypeptide N-acetylgalactosaminyltransferases (GalNAc-Ts). There are 20 different GalNAc-Ts in humans, each with a distinct substrate specificity and expression pattern. The GalNAc residue is added to the hydroxyl group of the serine or threonine residue in a reaction that requires UDP-GalNAc as the sugar donor (1).

2 Elongation of O-glycans: The second step in O-glycosylation involves the addition of additional sugar residues to the GalNAc residue. This step is catalyzed by a family of enzymes known as glycosyltransferases. The specific glycosyltransferases involved in O-glycan elongation vary depending on the tissue and the protein being modified. Some of the most common O-glycan structures include the core 1, core 2, and sialylated O-glycans (2).

3 Modification of O-glycans: The final step in O-glycosylation involves the modification of the O-glycan structures by various enzymes. These modifications include the addition of sulfates, fucoses, and sialic acids. The addition of these modifications can alter the biological activity of the protein and regulate its function. The specific modifications that occur on O-glycans depend on the tissue and the protein being modified (3).

In summary, the biosynthetic pathway leading to O-glycosylation in proteins involves the initiation of O-glycosylation by GalNAc-Ts, elongation of O-glycans by glycosyltransferases, and modification of O-glycans by various enzymes. The precise mechanisms and enzymes involved in each step of the pathway have been the subject of extensive research over the years and continue to be an active area of investigation.

References:

1 Ten Hagen, K. G. & Haltiwanger, R. S. Glycobiology DOI: 10.1093/glycob/cwx085 (2017).

2 Ju, T. & Cummings, R. D. Protein glycosylation: Challenging the dogma. Glycobiology DOI: 10.1093/glycob/cwu011 (2014).

3 Varki, A. Biological roles of glycans. Glycobiology DOI: 10.1093/glycob/cwt003 (2017).

**Notes of the Authors:** The ouput in this case appears largely correct, probably because, unlike in the previous example, it does not get in any specific detail of the process. Overall the description is superficial and does not go into any detail, as in the previous case, of the glycans precursor and terminal sequences. The references are again largely fabricated, except for the last one, where all details correspond to a highly cited review in the field, except for the DOI, which corresponds to a different article.

2. Here ChatGPT stalled [549 words] and we asked if it could complete the conclusions section, which it did with the text that follows.
